# Rapid local and systemic jasmonate signalling drives initiation and establishment of plant systemic immunity

**DOI:** 10.1101/2023.05.22.541689

**Authors:** Trupti Gaikwad, Susan Breen, Emily Breeze, Rana M. Fraz Hussain, Satish Kulasekaran, Marta de Torres-Zabala, David Horsell, Lorenzo Frigerio, Murray Grant

## Abstract

Successful recognition of pathogen effectors by plant disease resistance proteins (effector triggered immunity, ETI) contains the invading pathogen through a localized hypersensitive response (HR). In addition, ETI activates long-range signalling cascades that establish broad spectrum systemic acquired resistance (SAR). Using a novel and sensitive reporter we have been able to image the spatio-temporal dynamics of SAR. We demonstrate that local ETI triggered SAR signal generation, followed by rapid propagation and establishment in systemic responding leaves, is dependent on both jasmonate biosynthesis and perception. Further, ETI initiates calcium- and jasmonate-dependent systemic surface electrical potentials, reminiscent of those activated by herbivory but with slower propagation kinetics. Thus, jasmonate signalling is crucial to the initiation and establishment of systemic defence responses against a diverse range of phytopathogens.

## Introduction

Despite its discovery more than a century ago, our knowledge of the signalling processes underlying establishment, propagation and particularly initiation of plant systemic acquired resistance (SAR) remains fragmentary (1). Classically, SAR is established following effector triggered immunity (ETI) leading to the hypersensitive response (HR). SAR has also been reported to be activated via PAMP (Pathogen-Associated Molecular Pattern) recognition and virulent bacterial phytopathogens, although the latter has also been reported to trigger systemic induced susceptibility (2, 3).

Initially, salicylic acid (SA) and its volatile derivative methyl salicylate (MeSA) were proposed as key mobile signals (4-6). Today, many different molecules have been identified as possible SAR signals including azelaic acid (AzA) (7), glycerol-3-phosphate (G3P) (8), dehydroabietinal (9), nonanoic acid (NA) (10), pipecolic acid (Pip) (11), N-hydroxy-pipecolic acid (NHP) (12) with more recently, extracellular NAD(P] (13) and volatile monoterpenes α- and β-pinene (14) shown to be SAR inducing. These signalling molecules, their synthesis, activities and interactions have been extensively reviewed (1, 15-19).

Considering also that airborne defence cues activate SAR (20) has led to the consensus that SAR requires multiple signals, translocating apoplastically, symplastically and as volatiles to provide the necessary robustness to confer broad spectrum resistance against viral, bacterial, oomycete, fungal and insect pests with their diverse lifestyles and contrasting modes of infection (3).

The key SAR inducers such as SA and NHP are primarily synthesised *de novo*. Recently, single cell transcriptomics showed that during ETI genes in the NHP biosynthetic pathway (*FLAVIN*-*DEPENDENT MONOOXYGENASE 1, FMO1* and *AGD2-LIKE DEFENCE RESPONSE PROTEIN1, ALD1*) were strongly enriched in phloem companion cells implicating perception of a mobile inducing signal for NHP synthesis (21). Therefore, there must be primary inducing signals responsible for activating these bifurcate pathways given both branches are equally essential for SAR induction (16).

Early leaf detachment assays showed that SAR signal generation and translocation occur rapidly following primary leaf infection (22). Given ETI responses themselves require effector delivery, recognition and assembly of a resistosome complex (23, 24) it is most likely that the generation of the early inducing signals is intimately linked to an activated resistosome.

Both reactive oxygen species (ROS) and nitric oxide (NO) are integral to activation of SAR, via the oxidation of C18 unsaturated fatty acids on chloroplast lipids (3, 25). Indeed, AzA accumulates upstream of G3P but downstream of NO and H_2_O_2,_ being generated from the hydrolysis of C18 fatty acids released from the two nonionic lipid constituents of the thylakoid membrane, monogalactosyldiacylglycerol (MGDG) and digalactosyldiacylglycerol (DGDG) (26-28). Lipid signalling in SAR is also implied by the established role of lipid transfer proteins AZELAIC ACID INDUCED1 (AZI1) (29) and DEFECTIVE IN INDUCED RESISTANCE1 (DIR1) (26). Additionally, plants defective in SA, G3P, NO or ROS biosynthesis have reduced levels of Pip in distal tissues, underlining not only the complex metabolic interplay in establishment of SAR, but also the importance of the HR.

It is likely that enzymatic and/or non-enzymatic interactions underpinning the HR generate early SAR signals that can induce other signalling molecules. Recently, the rapid synthesis of two trans-acting small interfering RNAs (tasi-RNA), D7 and D8, from *TAS3a* have been shown to mediate SAR. These tasi-RNAs accumulated within 3 hours post infection (hpi) and cleaved the *Auxin response factors (ARF) 2, 3* and *4* to induce SAR. SAR is abolished by knockout of *TAS3a* or RNA silencing components required for tasi-RNA production, but critically, the levels of classical SAR signalling components were unaffected (30). Tasi-ARFs fit the key criteria for a SAR signal being rapidly transported (within 6 h of primary infection), induce resistance when locally applied and are essential for SAR. Genetic studies placed TAS3a downstream of G3P but still require distal SA production.

To elucidate early SAR signals it is essential to have a well characterised pathosystem. Recognition of *Pseudomonas syringae* pv. *tomato* DC3000 (DC) carrying *avrRpm1* by the RPM1 (Resistance to *Pseudomonas maculicola 1*) disease resistance protein (DC*avrRpm1*) (31) provides a robust ETI model to dissect signal initiation and transduction dynamics underlying SAR. RPM1 activation triggers early increases in cytosolic calcium (Ca^2+^_cyt_), beginning ∼1.5-2 hpi (32, 33) followed by lipid oxidation-triggered biophoton generation ∼3 hpi (34, 35) and visible leaf wilting ∼6 hpi. We have previously demonstrated RPM1 activation elicits rapid transcriptional reprogramming 4 hpi in systemic leaves, a large component mirroring the jasmonate-triggered systemic wound responses (36). Here we develop a novel jasmonate responsive SAR reporter which reveals unexpectedly rapid temporal and spatial transcriptional dynamics underpinning SAR. We show that establishment of SAR involves enzymatic production of a local jasmonate signal that symplastically propagates via the vasculature and symplastically via epidermal cells to systemic leaves, and is coupled to calcium and jasmonate dependent systemic surface electrical potentials.

## Results

### *JISS1* expression reveals temporal and spatial dynamics of early effector-R gene interactions

To monitor SAR transcriptional dynamics, we fused the promoter of *JASMONATE-INDUCED SYSTEMIC SIGNAL 1* (*JISS1*, At5g56980; previously known as *A70* (36)), a gene of unknown function annotated as induced by PAMPs (Araport11) and previously identified as an early SAR marker (36), to luciferase containing a DEAD box domain to ensure reporter turnover (Fig. S1). Homozygous *JISS1* promoter:*luciferase* (*JISS1:LUC*) lines showed rapid systemic luciferase activity following challenge with DC*avrRpm1* but not virulent DC, nor the type III secretion deficient strain DC*hrpA*, which elicits PTI responses but does not multiply significantly *in planta* (37), nor mock challenge (Fig. 1A). SAR signal propagation was remarkably rapid, with strong luciferase activity evident in the petiole of the challenged leaf ∼3 hpi (Fig. 1B). Within 30 min *JISS1:LUC* activity was established in leaves with two successive vasculature connections (38, 39), spreading to adjacent leaves within ∼50 min (Fig. 1B). By 4.5 hpi, 1 h prior to visible HR leaf collapse, systemic luciferase activity reached maximal intensity.

**Figure 1:**
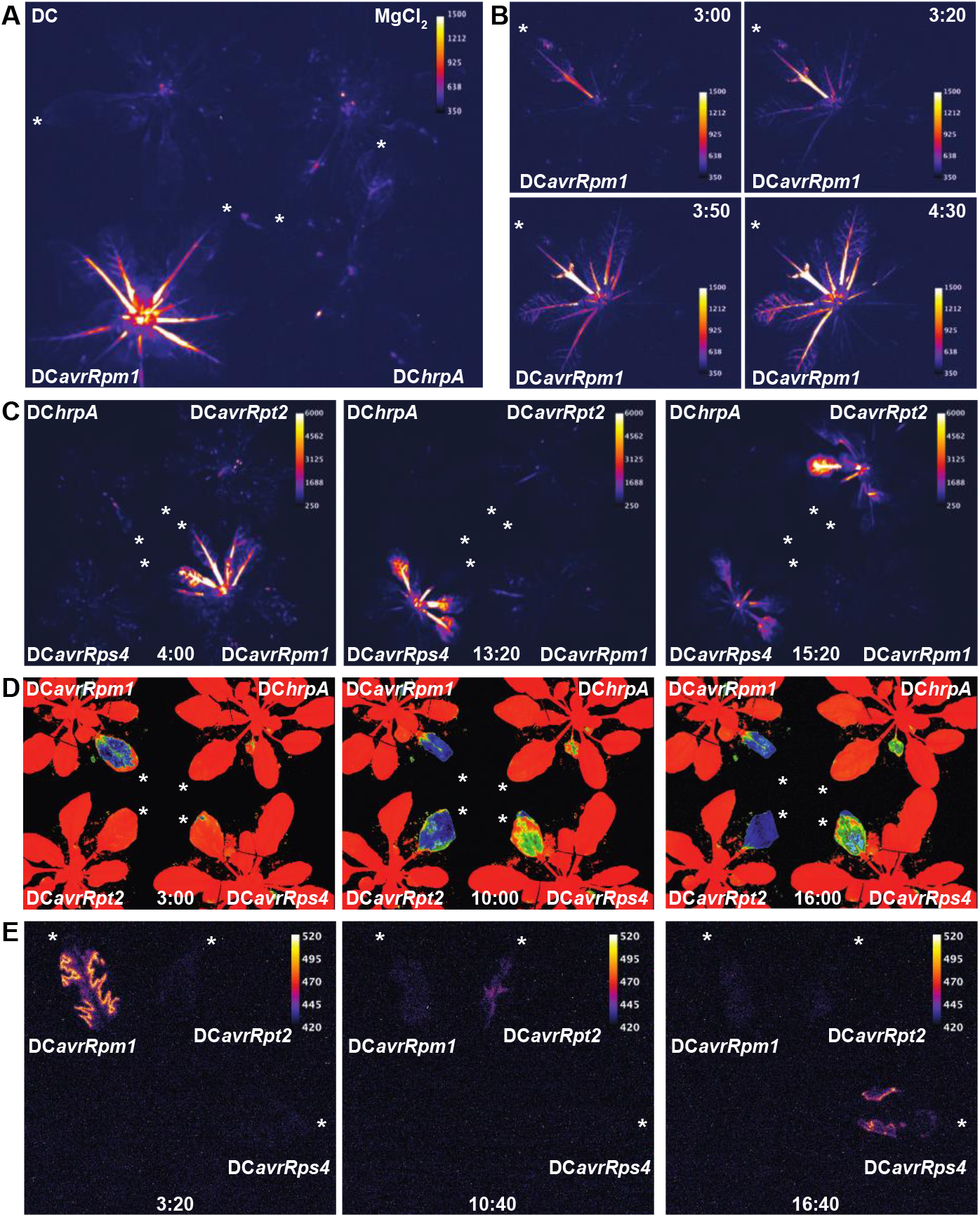
*JISS1* expression is induced systemically by ETI. White asterisk indicates infiltrated leaf and images are false coloured by signal intensity, as indicated by individual calibration bars. **(A)** Luciferase activity in *JISS1:LUC* plants following DC*avrRpm1*, DC, DC*hrpA* or mock (MgCl_2_) challenges at 4:30 hpi. Temporal spatial dynamics of luciferase activity in *JISS1:LUC* plants following DC*avrRpm1* challenge, initiating at 3 hpi. 3.20 hpi, 3.50 hpi and 4.30 hpi images capture the systemic spread of the signal over time. **(C)** Different *avr* genes display temporal specificity in activation of systemic *JISS1:LUC*; DC*avrRpm1* (4 hpi), DC*avrRps4* (13:20 hpi) and DC3*avrRpt2* (15:20 hpi), with a DC*hrpA* challenged control. **(D)** *F*_*v*_*/F*_*m*_ is strongly suppressed during ETI following DC*avrRpm1* (3 hpi), DC*avrRpt2* (10:00 hpi) or DC*avrRps4* (16:00 hpi) challenge. Orange indicates *F*_*v*_*/F*_*m*_ of healthy leaf (0.82), green represents *F*_*v*_*/F*_*m*_ of a compromised leaf (0.6), blue represents severely compromised *F*_*v*_*/F*_*m*_ of 0.32. **(E)** Biophoton generation is strongly induced during ETI following DC*avrRpm1* (3.20 hpi), DC*avrRpt2* (10:40 hpi) or DC*avrRps4* (16:40 hpi) challenge.

Similarly, challenge of *JISS1:LUC* leaves with DC*avrRpt2* or DC*avrRps4* also induced systemic luciferase activity following recognition by their respective resistance (R) proteins RPS2 (resistance to *Pseudomonas syringae* 2) (40) and RPS4 (resistance to *Pseudomonas syringae* 4) (41) (Fig. 1C). However, while the spatial pattern of reporter activity was identical for all three effectors, temporal luciferase dynamics were unique to each R protein (Fig. 1C and Mov. S1). To establish the temporal context of *JISS1* activation with respect to R protein activation we further investigated ETI dynamics using other non-destructive physiological readouts. The chloroplast acts as a sensor of biotic stress, best exemplified by a decrease in the quantum efficiency of photosystem II (*F*_*v*_*/F*_*m*_), which decreases markedly when leaves are infected with virulent DC, but not DC*hrpA* (42, 43) with earlier and more dramatic suppression elicited by ETI (44). Chlorophyll fluorescence imaging parameters associated with electron transport during photosynthesis were determined following challenge with DC*avrRpm1*, DC*avrRpt2*, DC*avrRps4* or DC*hrpA* (Fig. 1D). In addition, biophoton generation, which is indicative of chloroplast lipid peroxidation (35) and associated with initiation of the hypersensitive cell death (Bennett et al. 2005) was also assayed (Fig 1E). Collectively, these results revealed that timing of systemic signal initiation for these three R protein interactions was preceded in the local challenged leaf by significant disruption in chloroplast integrity and lipid peroxidation.

### Jasmonates are involved in ETI-induced systemic signalling events

To examine the role of JISS1 in SAR we crossed the *JISS1:LUC* reporter into classical SAR mutant lines: *npr1* (*NONEXPRESSOR OF PATHOGENESIS-RELATED 1*, a repressor of ETI but important for SAR (45-47); *npr1/3/4* (45), which is impaired in SA signal transduction but *npr3/4* are required for full RPS2 ETI (48); *nac19/55/*72 (altered regulation of SA accumulation) (49) and *sid2* (SA INDUCTION DEFICIENT 2; deficient in the accumulation of SA) (50). To facilitate crossing we identified the *JISS1:LUC* T-DNA insertion position (Fig. S1B). Surprisingly, when challenged with DC*avrRpm1*, all SAR compromised lines showed wild-type reporter dynamics, suggesting that *JISS1* is induced upstream of these key SAR components (Fig. 2A, Fig. S2). Furthermore, infiltration of known SAR elicitors, AzA, NA, Pip and NHP did not significantly increase local *JISS1:LUC* activity (Fig. 2B) and *JISS1:LUC* activity in an *fmo1* mutant, which is compromised in NHP production (12), was also wild-type in response to DC*avrRpm1* (Fig. 2C). Together, these data indicate that JISS1 accumulation occurs upstream and/or in parallel with previously characterized SAR elicitors.

**Figure 2:**
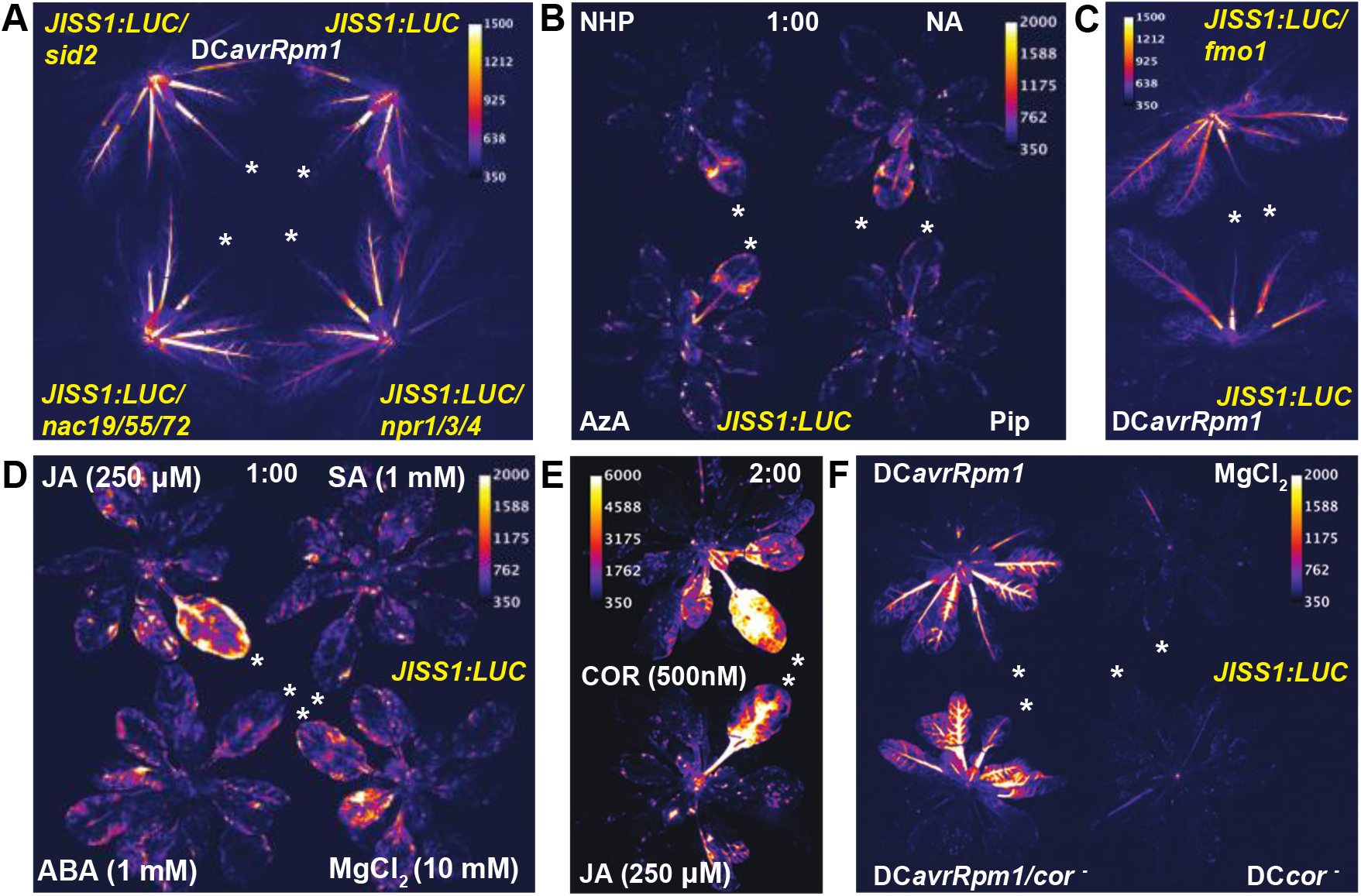
*JISS1:LUC* is activated by the jasmonate signalling pathway but not classical SAR elicitors. White asterisk indicates infiltrated leaf and images are false coloured by signal intensity, as indicated by individual calibration bars. **(A)** The classical SAR mutants, *sid2* (top left), *nac19/55/72* (bottom left) and *npr1/3/4* (bottom right) do not alter *JISS1:LUC* luciferase signatures following DC*avrRpm1* challenge (4 hpi). **(B)** Pipicolic (Pip), N-Hydroxypipicolic (NHP), nonanoic (NA) or azelaic (AzA) acids infiltrated (1 mM) into leaves of *JISS1:LUC* plants (1 hpi). **(C)** No attenuation of luciferase activity is observed in an *JISS1:LUC/fmo1* mutant line following DC*avrRpm1* challenge (4:30 hpi). **(D)** JA (250 µM) induces local *JISS1:LUC* signal propagation which is absent in leaves infiltrated with salicylic acid (SA, 1 mM), abscisic acid (ABA, 1 mM) or 10 mM MgCl_2_ (mock) (1 hpi). **(E)** Luciferase activity in *JISS1:LUC* leaves infiltrated with 250 µM JA or coronatine (COR, 500 nM) (1 hpi). **(F)** *JISS1:LUC* activity following challenge with DC or the coronatine deficient DC mutant DB4G3 (*cor*^*-*^) with or without *avrRpm1*. DC*avrRpm1cor*^*-*^ induced comparable *JISS1:LUC* systemic activity to DC*avrRpm1*, whereas no luciferase activity was detected in DC*cor*^*-*^ challenged leaves (3:50 hpi).

Given this unexpected result we next tested whether known key immunity associated phytohormones induced *JISS1*. Leaves infiltrated with SA (1 mM) or abscisic acid (ABA; 1 mM) also failed to elicit *JISS1:LUC* activity, however, luciferase signal was induced locally with jasmonic acid (JA; 250 µM), a core elicitor of systemic wound signalling (Fig. 2D). The jasmonates comprise JA and its derivatives, of which jasmonyl-isoleucine (JA-Ile) has been shown to have the highest binding affinity for the jasmonate co-receptor, the Skp/Cullin/F-box SCF^COI1^ (CORONATINE-INSENSITIVE PROTEIN 1)-JAZ1 E3 ubiquitin ligase complex (51-53). As part of its virulence strategy, DC produces the phytotoxin (and JA-Ile mimic) coronatine (COR) ∼10 hpi (52, 54). While 500 nM of COR strongly induced *JISS1:LUC* locally within 2 h (Fig. 2E) so did a DC*cor*^*-*^ mutant (55) expressing *avrRpm1* (Fig. 2F) ruling out COR as the elicitor.

### ETI elicits a rapid and propagative jasmonate signal essential for effective SAR

To further investigate the involvement of jasmonates in SAR, we investigated whether jasmonate biosynthesis and perception is important for SAR signalling. We phenotyped the JA biosynthetic mutant *aos* (ALLENE OXIDE SYNTHASE) (56) and *coi1-16*, which is defective in JA perception (57), for HR following challenge with DC*avrRpm1*, DC*avrRpt2*, DC*avrRps4* (Fig. S3A). In accordance with the differing temporal dynamics observed for each R protein in our luciferase assay (Fig. 1C and Mov. S1), collapse of the challenged wild type Col-0 leaf occurred first for DC*avrRpm1* (∼5.5 hpi), followed by DC*avrRpt2* (∼14 hpi) and then DC*avrRps4* (∼18 hpi) upon specific gene-for-gene recognition. Crucially, we observed comparable temporal responses to that of Col-0 in both the *aos* and *coi1-16* mutants (Fig. S3A), indicating that loss of jasmonate biosynthesis or perception does not abolish or attenuate ETI-induced local HR at high inoculum.

There is significant transcriptional overlap between wounding and SAR responses 4 hpi with DC*avrRpm1* (36). Re-examination of these data for *JASMONATE ZIM DOMAIN* (*JAZ*) genes, which act to reimpose repression of jasmonate signalling (52, 58), identified 9 of the 12 family members to be differentially expressed, including *JAZ10* (Table S1). A *JAZ10:GUS* reporter, induced systemically by local wounding (59), also shows remarkably similar spatial expression pattern in systemic leaves following DC*avrRpm1*, but not DC or DC*hrpA* challenges (Fig. 3A). However, systemic *JAZ10:GUS* expression is abolished in the jasmonate receptor mutant *coi1-16* (Fig. 3A) (57), further evidence that reporter activation is jasmonate dependent. We next crossed *JISS1:LUC* into the *coi1-16* and the JA biosynthetic mutant *aos1*. Distal expression of *JISS1:LUC* was not detected in either the *aos* nor *coi1-16* mutant backgrounds following DC*avrRpm1* challenge (Fig. 3B), and local application of JA or COR to J*ISS1:LUC*/*coi1-16* did not elicit a systemic signal (Fig. 3C). Additionally, RT-PCR of *JISS1* in systemic leaves of *aos* following DC*avrRpm1* challenge shows that expression of *JISS1* is reduced compared to Col-0 (Fig. S4B). Together, these data are consistent with a direct dependency on jasmonates for SAR but not for the local initiating HR.

**Figure 3:**
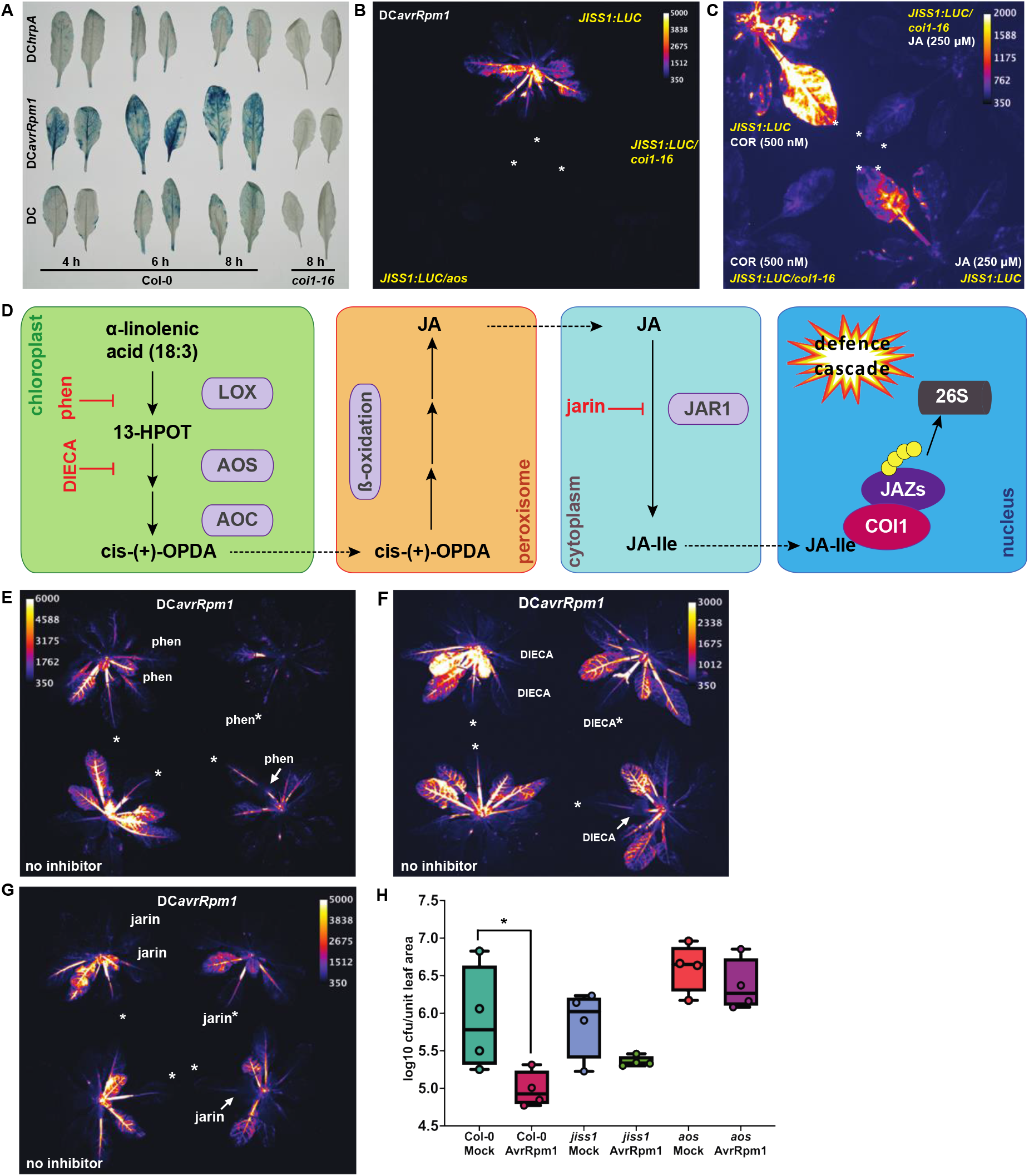
*JISS1:LUC* signal propagation is dependent on JA biosynthesis and perception. White asterisk indicates infiltrated leaf and images are false coloured by signal intensity, as indicated by individual calibration bars. **(A)** GUS activity in representative systemic leaves of a *JAZ10:GUS* reporter line at times indicated compared to a *JAZ10:GUS/coi1-16* line 8 hpi. **(B)** *JISS1* is not induced in DC*avrRpm1-*treated *JISS1:LUC/aos* nor *JISS1:LUC/coi1-16* lines (5 hpi) **(C)** Luciferase activity is absent in an *JISS1:LUC/coi1-16* mutant but not *JISS1:LUC* leaves following infiltration with 250 µM JA or 500 nM of the JA mimic coronatine COR) (1 hpi). **(D)** Schematic of JA biosynthetic pathway highlighting positions of JA inhibitor activity. **(E-G)** Treatment with phenidone (2 mM; 4:40 hpi) **(E**), DIECA (2.5 mM; 5:30 hpi) **(F)** or Jarin-1 (25 µM; 5:10 hpi) **(G)** inhibits DC*avrRpm1*-induced *JISS1:LUC* activity. Key to **(E-G)**, top left: two systemic leaves on right of local leaf pre-infiltrated with inhibitor. Top right: local leaf co-infiltrated with inhibitor. Bottom right: petiole of immunised leaf treated with inhibitor. Bottom left: no inhibitor. **(H)** SAR growth curve of *Psm4* following DC*avrRpm1* or mock immunising challenge on Col-0, *jiss1* and *aos* mutants. Error bars, mean ± SE (n=4), statistical significance p=0.0653 (Students’ t-test).

Aside from JA-Ile, many jasmonate molecules have been demonstrated to have biological activity *in planta*, including OPDA (12-oxophytodienoic acid), although OPDA is not perceived by the SCF^COI1^ receptor complex (60). We tested the JA biosynthetic inhibitors phenidone, a lipoxygenase inhibitor (61), diethyldithiocarbamic acid (DIECA) which interferes with octadecanoid signalling (62) and jarin-1, an inhibitor of JA-Ile synthetase (63) (Fig. 3D). Concurrent DC*avrRpm1* challenge and treatment with phenidone (2 mM), DIECA (2.5 mM) or jarin (25 µM) via infiltration into the adjacent systemic responding leaves, co-infiltration or petiole application, abolished, or markedly attenuated, DC*avrRpm1-*elicited *JISS1:LUC* activity (Fig. 3E-G). These data reinforce our finding that jasmonates play a key role in SAR, both in signal generation and initiation in naïve responding leaves and JA-Ile biosynthesis appears to be necessary to elaborate effective signal transduction.

The *coi1* mutant is known to be more resistant to local DC challenge than wild-type Arabidopsis (64). We assessed SAR to virulent *P. syringae* pv. *maculicola* race 4 (*Psm4*) in the *aos* and *coi1-16* mutants alongside a *JISS1* T-DNA insertion loss-of-function line (*jiss1*, Fig S4B, C). SAR assays in *jiss1* were inconsistent. Although enumeration of bacterial counts of *Psm4* in systemic leaves of *jiss1* following DC*avrRpm1* challenge were not significantly different (p>0.05) from those observed following mock immunizing challenge in replicated SAR assays, they were consistently lower (Fig. 3H; p=0.0653, df=6). Consequently, we cannot conclusively state that SAR is abolished in this line. Moreover, two homologues of JISS1 exist in Arabidopsis, AT4G26130 and AT2G26110 with 56% and 35% amino acid similarity, respectively. Thus, it is possible functional redundancy may exist in JISS1 function, accounting for our variable SAR assays. Crucially, neither jasmonate mutant exhibited SAR (Fig. 3H and Fig. S3B). In addition, *aos* showed a trend towards increased susceptibility relative to that of Col-0 across multiple independent assays.

Collectively these data imply that ETI elicits rapid *de novo* synthesis of a jasmonate signal that propagates systemically and is essential for effective establishment of SAR.

### JISS1 signal predominately localizes to the vasculature and epidermal endoplasmic reticulum

To further understand the role of JISS1 in SAR we investigated its localization and expression following ETI elicitation. Since JISS1 was previously annotated as chloroplast localized (TAIR10; AtSubP), we generated a truncated JISS1-GFP fusion encoding the first 89 amino acids of JISS1 (*JISS1*_pro_: JISS1^1-89^-GFP), to include its predicted chloroplast transit peptide. GFP signal was detected within 4.5 hpi across the entire systemic leaf (Fig. 4A), predominately in the vasculature and, interestingly, epidermal cells of both the laminar (Fig. 4B) and petiole (Fig. 4C, Fig. S5A) of the systemic responding leaf. Unexpectedly, the fusion protein localized to the endoplasmic reticulum (ER) in leaf epidermal pavement cells (Fig. 4D), suggestive of a SAR signal moving symplastically, consistent with the abolition of SAR in Arabidopsis lines overexpressing PLASMODESMATA-LOCATED PROTEIN 5 in which plasmodesmata permeability is significantly reduced (65). As observed with *JISS1:LUC* activity, GFP expression in systemic leaves was markedly suppressed by pre-treatment with either phenidone or DIECA, JA biosynthetic inhibitors (Fig. S5B). To confirm JISS1 localization we generated a full-length JISS1-GFP reporter line (*JISS1*_pro_:JISS1-GFP) which showed identical subcellular localization to *JISS1*_pro_:JISS1^1-89^-GFP (Fig 4D, Fig S5C).

**Figure 4:**
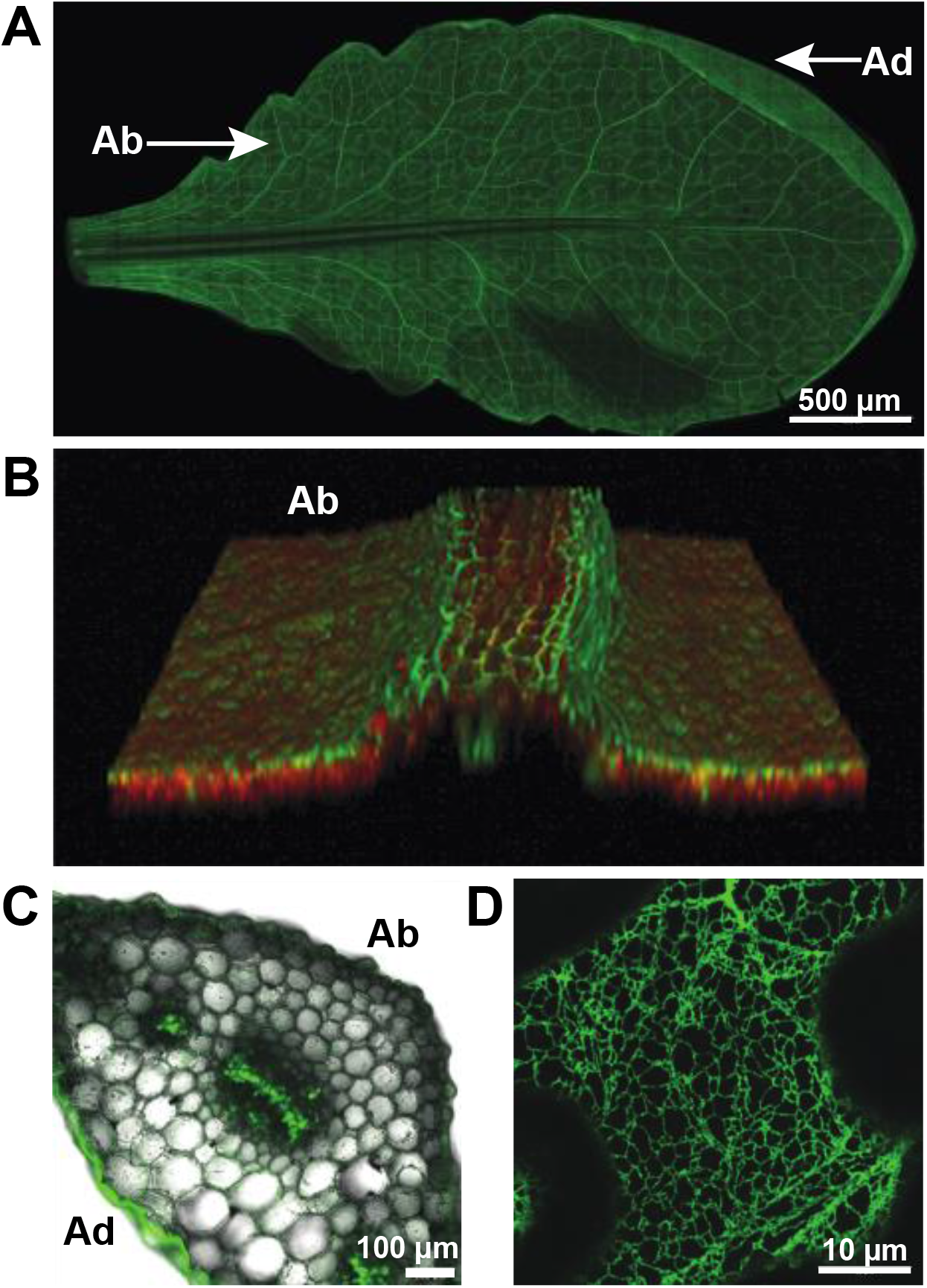
JISS1 signal propagates symplastically through the epidermis and vasculature and localises to the endoplasmic reticulum. Confocal images of global *JISS1:*JISS1^1-89^-GFP expression in representative systemic leaves following DC*avrRpm1* challenge. Ab, abaxial surface; Ad, adaxial surface. **(A)** JISS1^1-89^-GFP expression showing JISS1-GFP accumulates throughout the leaf, with the signal predominantly in the vasculature. Scale bar, 500µm; **(B)** Still from Supp Movie 2 of JISS1^1-89^-GFP highlighting that the GFP signal is predominately restricted to the central vein and epidermal cell layer. JISS1^1-89^-GFP expression in vasculature of petiole and epidermal cell layer. Scale bar, 100 µm. **(D)** Subcellular localisation of JISS1^1-89^-GFP is restricted to the endoplasmic reticulum of epidermal cells. Scale bar, 10 µm.

### Systemic electric signal propagation is a general feature of ETI activation

Wound-induced accumulation of JA and JA-Ile in undamaged distal leaves together with the altered expression of jasmonate-responsive genes, is preceded by rapid generation of wound-activated surface electrical potentials (WASPs), caused by plasma membrane depolarization (59). WASPs are mediated by glutamate triggered systemic activation of cytosolic Ca^2+^ (Ca^2+^_cyt_) signalling via specific vasculature localized members of the glutamate-like receptor family, GLR3.3 and GLR3.6 (59, 66), cation-permeable ion channels, which also contribute to systemic defences to herbivory (67). Given the comparable propagative response we observed with *JISS1:LUC*, we examined whether electrical surface potentials contribute to ETI mediated SAR signalling. Electrodes were attached to the midrib/petiole junction of the local challenged leaf (‘infiltrated’) and fully expanded leaves immediately adjacent (‘adjacent’) and at ∼180° to the challenged leaf (‘distal’) and changes in leaf surface potentials recorded (Fig 5A).

**Figure 5:**
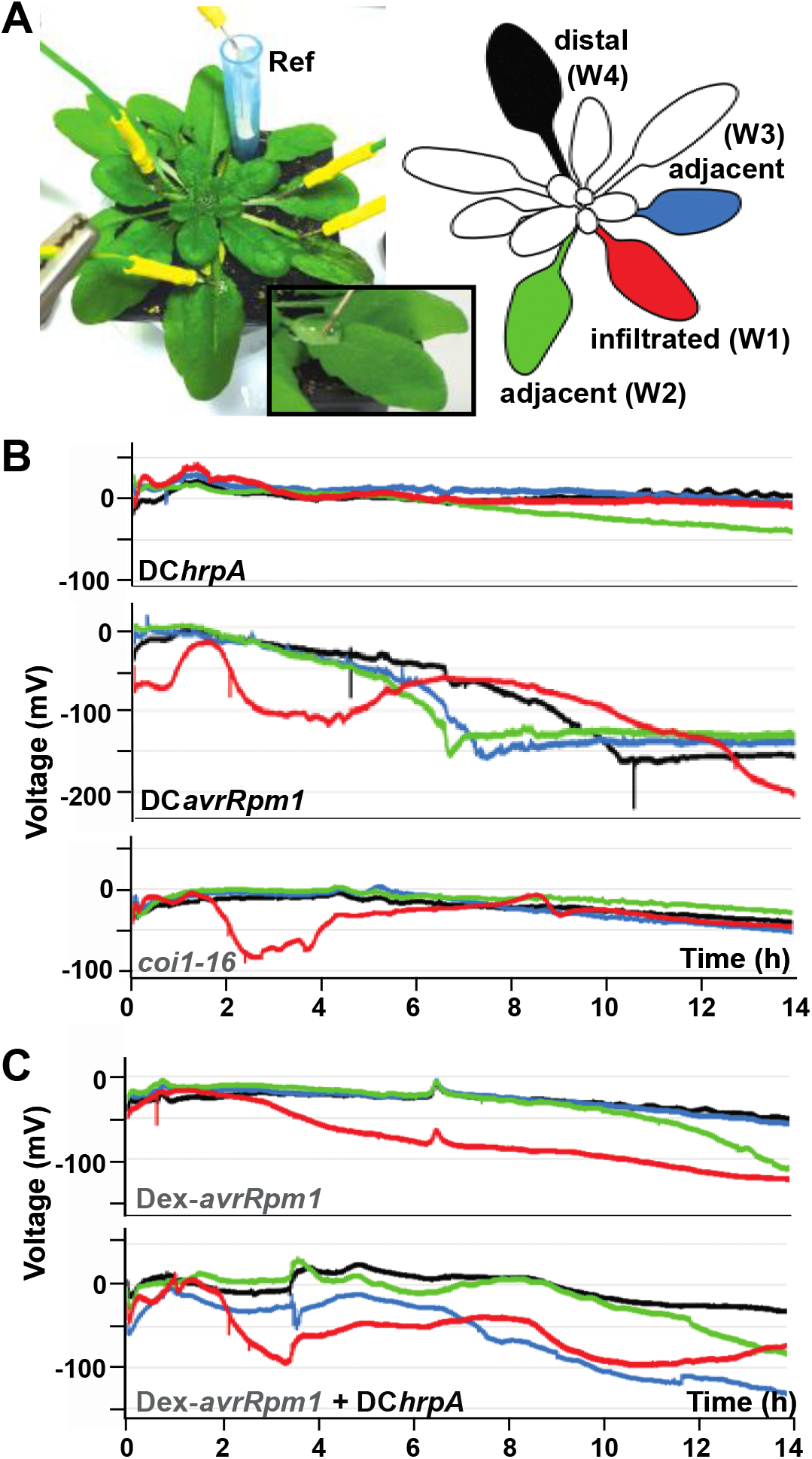
Systemic electric signal propagation is a general feature of ETI activation. **(A)** Schematic of plant electrophysiology experimental set-up with cartoon showing spatial sampling and colour coding of leaves for working electrodes (W). Inset illustrates electrode positioning. Ref, reference electrode. **(B)** DC*avrRpm1* but not DC*hrpA* challenged leaves (red) show an initial depolarisation ∼2 hpi and subsequent repolarisation. From 4-7 hpi SISPs are propagated in the two systemic leaves (blue and green) adjacent to the DC*avrRpm1* immunized leaf, with the distal leaf (black) responding later (from ∼7 hpi). No SISPs are observed in the DC*hrpA* treatments. c*oi1-16* mutant shows depolarisation of the DC*avrRpm1* challenged leaf but cannot initiate SISPs. **(C)** Dex-induced *avrRpm1* expression does not replicate DC*avrRpm1* induced SISPs, however, infiltration of DC*hrpA* 1 h after Dex application re-instigates SISPs.

Challenge with DC*hrpA* did not induce specific changes in leaf surface potentials. Challenge with DC induced a small depolarization of the local challenged leaf only but no SIPSs (Fig. 5B, Fig S6A). In contrast, challenge with DC*avrRpm1* induced Systemic Immunity Surface Potentials [SISPs]) in both the local and systemic leaves (Fig. 5B). Unlike WASPs and herbivory responses which travel at speeds well in excess of millimetres per second (68), SISPs were observed to be slower variation potentials (measured over several hours), which largely mirror the spatial dynamics of systemic *JISS1:LUC* activity. Compared to DC inoculation, the DC*avrRpm1*-challenged leaf (red trace) consistently showed a depolarization of ∼-100 mV of duration ∼2 h (Fig. 5B), the initiation of which strongly correlated to prior biophoton generation, suppression of *F*_*v*_*/F*_*m*_ and *JISS1:LUC* systemic signalling (Fig. 1). Following repolarization of the challenged leaf, SISPs were detected in adjacent leaves (blue and green traces) with maximal depolarization ∼7 hpi. SISPs were also detected in distal leaves (black) but maximal depolarization was not achieved until ∼10 hpi (Fig. 5B). As both DC*avrRpt2* and DC*avrRps4* also trigger SISPs (Fig. S6A) with respective timings largely consistent with initiation of suppression of *F*_*v*_*/F*_*m*_ (Fig. 1D), we conclude SISPs are specifically elicited by ETI.

Given our previous finding that ETI-induced systemic signalling requires effective jasmonate perception, we further investigated SISP generation in the *coi1-16* mutant. Only the DC*avrRpm1* challenged leaf underwent depolarization, SISPs were not detected (Fig. 5B). Surprisingly, despite eliciting a local visible HR, SISPs were not induced by conditional (dexamethasone, DEX) induction of *avrRpm1*. Instead, a steady depolarization of the induced leaf with no repolarization was observed (Fig. 5C). However, co-infiltration of DEX-induced leaves with DC*hrpA* (Fig. 5C) or DC (Fig. S6B) broadly recapitulated SISP changes, implying activation of PTI is required for SISP generation.

We also measured SISPs in the *glr* mutants (*glr3*.*3, glr3*.*6*, and *glr3*.*3/glr3*.*6*) whose functions are necessary for full wound and herbivory responses (59, 67) and in the *jiss1* mutant. Mirroring WASP attenuation (59), SISPs were totally abolished in the double *glr3*.*3/glr3*.*6* mutant (Fig. 6A) and largely abolished in the single *glr3*.*3* and *glr3*.*6* mutants, with *glr3*.*6* only showing depolarization of the local immunized leaf (Fig. 6A) (67). (34). As for *coi1-16*, no SISPs were observed in *jiss1* (Fig. 6A).

**Figure 6:**
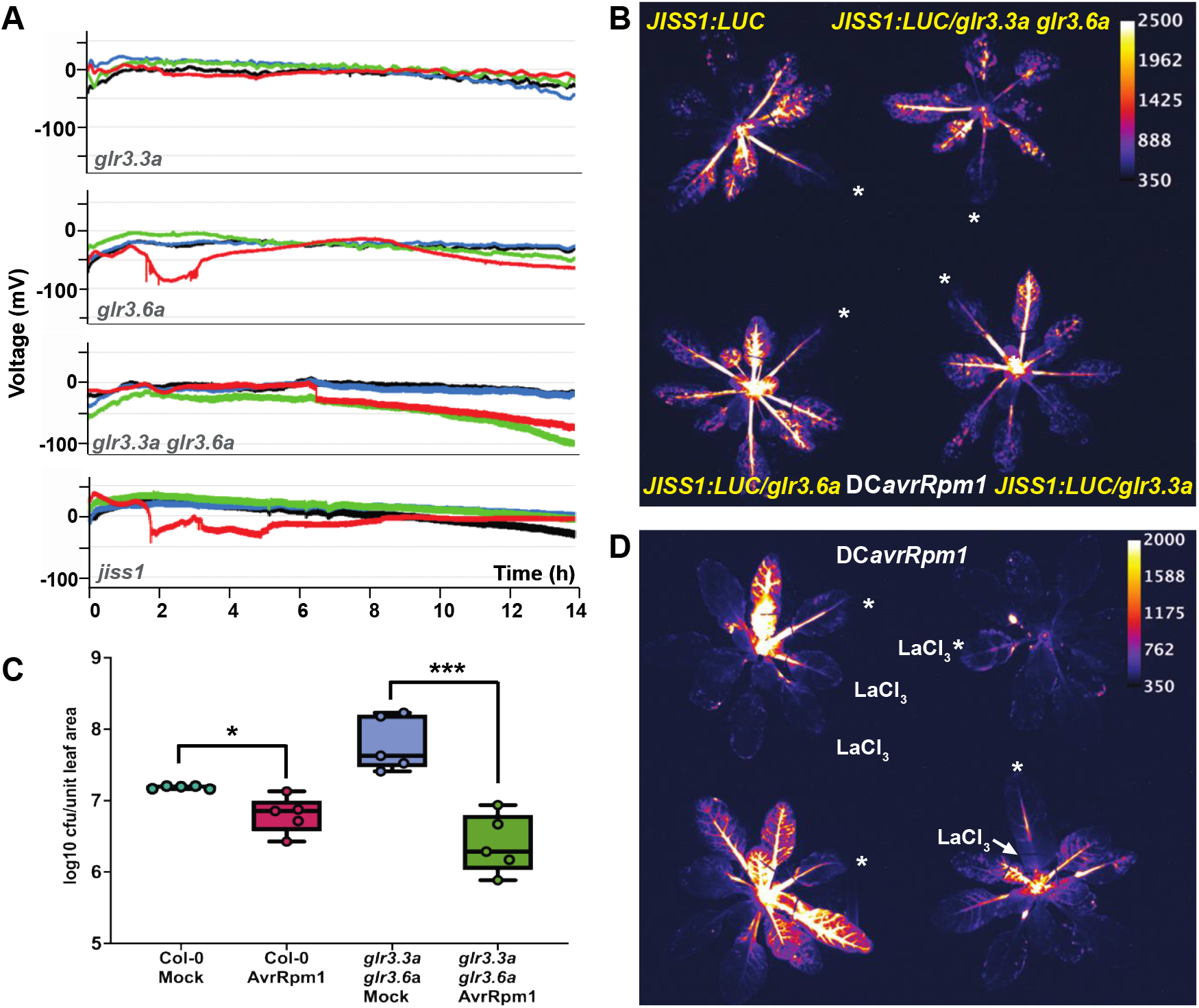
JISS1 systemic signal is calcium dependent. **(A)** Glutamate receptor mutants *glr3*.*3a, glr3*.*6a* and *glr3*.*3a glr3*.*6a* do not show development of SISPs following DC*avrRpm1* challenge, although depolarisation of the infiltrated leaf occurs in *glr3*.*6a*. Additionally, *jiss1* shows depolarisation of the DC*avrRpm1* infiltrated leaf but no development of SISPs. **(B)** DC*avrRpm1* challenged *JISS1:LUC/glr3*.*3a glr3*.*6a* mutant (top right), *JISS1:LUC*/*glr3*.*3a* (bottom right) and *JISS1:LUC/glr3*.*6a* (bottom left) all exhibited comparable systemic luciferase activity to that of *JISS1:LUC* (top left) (4:10 hpi). White asterisk indicates infiltrated leaf. Luciferase images are false coloured by signal intensity, as indicated by calibration bar. **(C)** SAR growth curve of *Psm4* following DC*avrRpm1* or mock pre-treatment on wild-type (Col-0) and *glr3*.*3a glr3*.*6a* mutant. Error bars represent the mean ± SE (n=5) and statistical significance (P<0.005) was determined by Student t-test. **(D)** Treatment with the calcium inhibitor LaCl_3_ (1 mM) inhibits DC*avrRpm1*-induced *JISS1:LUC* activity compared to control (lower left) (4:50 hpi). Top left: two systemic leaves on right of local leaf pre-infiltrated with inhibitor. Top right: local leaf co-infiltrated with inhibitor. Bottom right: petiole of immunised leaf treated with inhibitor. Bottom left: no inhibitor. White asterisk indicates infiltrated leaf and luciferase images are false coloured by signal intensity, as indicated by calibration bar.

### Calcium signalling is necessary for systemic immunity

Given the finding that ETI alone is not sufficient to induce SISPs, we further investigated the relationship between SISPs and SAR. To this end we crossed *JISS1:LUC* into the *glr3*.*3, glr3*.*6* and *glr3*.*3/glr3*.*6* mutants. Consistent with an intact SAR to *P. syringae, JISS1:LUC* dynamics were unaltered in all loss-of-function *glr* lines tested (Fig. 6B, C). These data suggest that *JISS1* signalling precedes, or is independent of, SISP generation and reinforces a core role for jasmonates and *JISS1* in specifying SAR to *P. syringae*.

In summary, AvrRpm1-RPM1 recognition initiates jasmonate-dependent *JISS1* systemic signalling and SISPs. SISPs are also dependent upon jasmonate signalling, requiring functional JISS1 and COI1, but vasculature-specific GLR Ca^2+^ channels which link systemic electrical signal propagation in response to wounding (59) and herbivory (67) do not abolish SAR to *Psm4* implying SISPs may encode another SAR signal, for example for herbivory responses. However, application of the calcium channel blocker LaCl_3_ (1 mM) to systemic leaves abolished DC*avrRpm1* induced *JISS1:LUC* activity, as did co-infiltration of LaCl_3_ with DC*avrRpm1* into the challenged leaf, with petiole application markedly attenuating *JISS1:LUC* activity (Fig. 6D). Therefore, Ca^2+^ signalling, which is intimately linked to resistosome activation (24), is required for effective SAR, implying involvement of other GLRs and/or alternative calcium ion channels.

## Discussion

SAR is broad spectrum and it is highly unlikely that a single primary mobile inducing signal can alone establish resistance to viral, fungal, oomycete and bacterial pathogens, and insect pests with their diverse lifestyles and virulence strategies. A variety of signalling molecules have been associated with SAR from the N-hydroxylated non-protein amino acid (NHP) and the hormone SA to volatile and non-volatile fatty acid derivatives. SA, NHP and these volatiles are synthesized *de novo* and thus their synthesis itself requires decoding of upstream inductive signal(s). For the volatiles, reactive species elicitation is the most harmonious explanation. Our current understanding of signal generation, translocation and establishment in systemic responding leaves is restricted by a lack of information on spatial temporal dynamics of SAR activation. Here we have developed and characterized a novel reporter that faithfully reports systemic signalling initiation and translocation from both coiled-coil (CC) NLRs (CNLs) and Toll/interleukin-1 receptor (TIR) NLRs (TNLs). Further, the dependency on a jasmonate signal led us to identify the propagation of HR mediated electrical surface potentials in systemic responding leaves.

The *JISS1:LUC* reporter was developed from a previous study (36) that identified systemic induced genes within 4 hpi with DC*avrRpm1*. Further transcriptomic analysis of systemic responding leaves revealed a significant over-representation of genes associated with jasmonate biosynthesis/regulation, wound and herbivory responses by 4 hpi, but not following DC challenge nor in *rpm1* or RPM1 signalling mutants. We established *JISS1:LUC* reporter dynamics during HR elicited by two CNLs (RPM1, RPS2) and a TNL (RPS4) using a combination of chlorophyll fluorescence (43, 44) and biophoton (34) imaging. *JISS1:LUC* reporter activation was specific to ETI, with no systemic activation following mock, PAMP activating DC*hrpA* or virulent (DC) challenges. Following RPM1 activation, induction was rapid and almost concomitant with biophoton generation and strong suppression of *F*_*v*_*/F*_*m*_ (Fig. 1) with signal translocation across systemic leaves within 50 min, the distribution of which is indicative of acropetal movement (38, 69). Systemic *JISS1:LUC* reporter activation by RPS2 and RPS4 followed biophoton initiation and *F*_*v*_*/F*_*m*_ activation. Interestingly, despite earlier *F*_*v*_*/F*_*m*_ suppression and biophoton induction, the systemic signal activation was faster in RPS4 than RPS2, though the strength and spatial distribution of the systemic signal was similar (Fig. 1). We hypothesize that these differences in SAR signal generation reflect differences in resistosome formation and activity between TNLs and CNLs (23, 24)

Notably, none of the key SAR suppressing mutants, *sid2, npr1/3/4, fmo1* or *nac19/55/72* affected systemic signal generation (Fig. 2). Similarly, infiltration of the SAR inducers NA, AzA, PA or NHP activated *JISS1:LUC*. Of the key hormones associated with plant immunity crosstalk, ABA and surprisingly, SA, failed to activate the *JISS1:LUC* reporter whereas JA did, albeit only locally with no systemic activation. Similarly, the DC produced bioactive JA-Ile mimic, coronatine, also activated *JISS1:*LUC, again only locally, but at a concentration 500-fold lower than JA. Although DC did not activate *JISS1:LUC*, we confirmed challenge with the coronatine deficient DC*avrRpm1* still gave identical *JISS1:LUC* temporal and spatial activation dynamics ruling out COR as the inductive signal.

We crossed the *JISS1:LUC* reporter into the F-box protein CORONATINE INSENSITIVE 1 (COI1) jasmonate receptor mutant *coi1*-16, part of a Skp/Cullin/F-box SCF^COI1^ E3 ubiquitin ligase complex (51-53), and into the ALLENE OXIDE SYNTHASE (*aos*) JA biosynthetic mutant (56) (Fig. 3D). Upon COI1 binding of JA-Ile, SCF^COI1^ recruits JASMONATE ZIM DOMAIN (JAZ) transcriptional repressors for proteasomal degradation (70), de-repressing *MYC2, MYC3* and *MYC4* encoding basic-helix-loop-helix transcription factors which activate JA responsive gene via their interaction with the Mediator complex subunit, MED25 (71). *JISS1:LUC* activity was totally abolished in both *coi1-16* and *aos* (Fig. 3B), indicating a crucial role for jasmonates in initial systemic signal generation and propagation. Neither COR nor JA infiltration activated luciferase in *coi1*-16 *JISS1:LUC* plants (Fig. 3C). While SAR was abolished in *coi1*-16 plants (Fig. S3B) and *coi1* mutants failed to activate *JAZ10* in systemic tissue (Fig. 3A), these results are confounded by the already heightened resistance of *coi1* to *P. syringae* due to elevated SA (72). We had previously shown (36) that SAR was compromised in the JA biosynthetic mutant, *opr3 (*12-OXOPHYTODIENOATE REDUCTASE 3) (73) and JA signalling mutant, *jin1* (JASMONATE-INSENSITIVE 1; *myc2*) (74), and here we showed that the *aos* mutant also failed to activate SAR. Consistently, application of three jasmonate biosynthesis inhibitors, jarin, DIECA or phenidione, significantly attenuated SAR signal generation, propagation and/or perception, reinforcing a role for jasmonates in SAR signalling.

It may seem counterintuitive that jasmonates underpin a primary SAR signal, given JA/SA antagonism in biotrophic immunity. Although not extensively referenced, increases in endogenous JA occur in parallel to SA accumulation during ETI (75), with Mur et al. (76) first reporting a link between SA/JA cross talk and reactive oxygen species. Indeed, the HR is widely thought to be triggered by singlet oxygen generation, leading to lipid peroxidation (35). Our data support the model that early ETI leads to chloroplast ROS generation captured by the strong suppression of *F*_*v*_*/F*_*m*_ (44), impacting chloroplast integrity and resulting in lipid peroxidation as illustrated by biophoton generation (34, 35). These events result in an early accumulation of both enzymatic and non-enzymatic chloroplast galactolipid derived oxylipins (77, 78) and underpin generation of a jasmonate based SAR signal and *JISS1:LUC* activation. As further evidence of a central role for JA in SAR, both fatty acid desaturase and chloroplast galactolipid mutants necessary for JA synthesis are SAR deficient (25, 27) and JA levels have been shown to increase 75 fold 5-10 hpi with DC*avrRpm1* (78).

Interestingly, JA dependency in ETI has been reported for the AvrRpt2-RPS2 interaction. JA signalling positively regulates the establishment of RPS2-mediated ETI. *Psm* ES432 *avrRpt2* challenge elicited a rapid activation of JA-responsive genes in challenged leaves with increases in JA (8 hpi) and JA-Ile (12 hpi). HR was, unexpectedly, dependent on SA and the SA receptors NPR3 and NPR4, but not on NPR1 which has been reported to repress ETI (45, 46). Expression of *JAZ1, JAZ10* and JA biosynthetic pathway *LOX3* genes were attenuated in both *npr3/npr4* and *sid2* mutants but exogeneous application of methyl jasmonate could rescue RPS2 mediated ETI (48). While COI1 was required for the initial activation of JA-responsive genes, ETI was diminished in *coi1* implying that the canonical JA pathway is required for subsequent signal amplification (48).

As we saw no alteration in dynamics of RPM1 induced *JISS1* reporter activity in *npr1, sid2*, the *npr1/3/4* triple mutant nor in *nac19/55/72* lines compromised in SA accumulation, we conclude that *JISS1* reports an independent ETI mediated systemic signalling pathway, though Lui et al’s (48) data may also explain the delay in *JISS1:LUC* mediated signalling seen between RPS2 and RPS4 challenges (Fig. 1). Spatial separation of JA and SA signalling in ETI would be a parsimonious explanation for the counterintuitive SA/JA dependency and the previously published (45, 79-81) but often neglected importance of short distance local cellular signalling in ETI responses. Indeed, intravital time-lapse imaging of the spatial distribution of SA and JA markers following RPS2 ETI elicitation revealed clear separation in distinct concentric and temporally developing domains (SA, inner/early and JA outer/later) between the ETI responding cells (82) elegantly explaining the apparent SA/ JA antagonism conflict observed (48) and the COI1 dependency of the *JISS1* reporter. A recent study (83) reinforces the need to consider spatial cellular context during ETI. Potato virus Y (PVY)- elicited local transduction of ROS signalling in chloroplasts of potato palisade mesophyll and upper epidermal cells was SA dependent. Pharmacological intervention abrogated these spatially distinct chloroplast redox changes which were underpinned by transcriptional reprogramming.

The *JISS1* reporter spatial pattern mirrored the jasmonate dependent wound associated surface potentials (WASPs) (59). Measurement of leaf surface variation potentials demonstrated that ETI triggered SAR resulted in the generation of SISPs, albeit the dynamics differed from systemic wound/herbivory signalling which occurs within minutes (68), whereas ETI systemic signal propagation across similar distances was in the order of tens of minutes. ETI responding leaves triggered large surface depolarizations (∼ 100 mV), at least similar to WASPs, and this depolarisation is necessary for activation of the JA pathway. Interestingly conditionally expression of *avrRpm1 in planta* was insufficient to recapitulate SISPs generated following avirulent pathogen challenges and required addition of PAMPs. Notably, wounding and herbivory release damage-associated molecular patterns (84, 85) and these results suggest that the recently described PTI/ETI potentiation (86) extends to SAR signalling. Both *coi1* and *jiss1* knockout lines only show depolarisation of the challenged but not systemic responding leaves.

The glutamate receptors, *glr3*.*3, glr3*.*6* and the *glr3*.*3/glr3*.*6* double mutant lines abolish WASPs and strongly attenuated expression of wound induced JA-response genes in systemic leaves (59, 66). Consistently, *glr3*.*3, glr3*.*6* and the *glr3*.*3/glr3*.*6* abolished SISPs with *glr3*.*3* exhibiting markedly less depolarisation of the challenged leaf (Fig. 6A, red trace) than *glr3*.*6*. Unexpectedly, none of the *glr* mutants tested abolished *JISS1:LUC* activity (though signal appeared attenuated in the *glr3*.*3* background) nor abolished SAR to *Psm4* (*glr3*.*3/glr3*.*6*), despite restricting herbivory (67). While the underlying mechanisms require further investigation, the parallels with wounding/herbivory responses are consistent with recruitment of similar signalling components, albeit generating different signatures and amplitudes. Thus, SISPs may represent another jasmonate dependent ETI generated systemic signal that are interpreted as a signature of herbivory, one of a suite of signals that collectively encode reprogramming of distal leaves to confer resistance to the diverse range of pathogens SAR successfully restricts.

Clearly more research is necessary to understand GLR mechanism of action in generation SISPs. In Arabidopsis, an apoplastic glutamate sensor detected increases in extracellular glutamate within wounded leaves (67). It is therefore possible that glutamate activates one or more of the GLRs. GLR3.3 and GLR3.6 localise to the leaf vasculature. Interestingly, GLR3.3 is primarily ER localised in phloem sieve elements whereas GLR3.6 was in xylem contact cell tonoplast membranes (66). Patch clamping showed GLR3.3 acted as a Ca^2+^ and Na^+^ non-selective channel (87). Applying the same experimental strategy as used with the jasmonate inhibitors we show that inhibition of Ca^2+^ channels also abolishes *JISS1:LUC* activity, thus both SISPs and *JISS1* activation require initial Ca^2+^ signalling. Interestingly, LOX6 is stimulated by increases in cytosolic Ca^2+^ and thus may be responsible for contributing to JA synthesis (88, 89).

In summary, there are remarkable parallels between systemic wound signalling and elicitation of SAR, implying recruitment of common signalling components. Systemic wound/herbivory signalling occurs within minutes (68), whereas ETI systemic signal propagation, initiated following local biophoton generation and suppression of *F*_*v*_*/F*_*m*_, tens of minutes. These ETI responses require jasmonate-derived signals and, like wounding, generate SISPs, which are both jasmonate dependent and require GLRs. We propose SISPs propagate the mobile information decoded systemically to initiate defences to herbivory (67), whereas rapid jasmonate induction establishes defence to biotrophs. Furthermore, JISS1, an ER-localised protein, functions as a dynamic SAR reporter capturing a propagative jasmonate signal rapidly moving from the immunized leaf through the vasculature and symplastically via epidermal cells to systemic responding leaves. This study lays the foundation for dissecting further the complex molecular mechanisms underpinning SAR.

## Materials and Methods

### Arabidopsis growth conditions

*Arabidopsis thaliana* were grown for 4-5 weeks in a compost mix (Levingston F2 compost) in a controlled environment growth chamber programmed for 10 h day (21°C; 120 μmol m^−2^ s^−1^) and 14 h night (21°C) regime with 60% relative humidity, as previously described (54, 90). Growth conditions for the SAR growth assays were 16 h day (21°C; 120 μmol m^−2^ s^−1^) and 8 h night (21°C) with relative humidity at ∼80%.

### Origin of transgenic Arabidopsis lines

The *JISS1:LUC* plants were generated as previously described (91). In brief *luc2P* was digested out of Promega pGL4-11 and cloned into pCAMBIA1302 via Kpn1 and Pml1 restriction sites creating pC1LUCP. A 1631 bp promoter fragment of JISS1 was PCR amplified from Col-0 genomic DNA using the primers:

JISS1F + KpnI AAT<u>CCATGG</u>TCAACCGTAAAAGGTCGGTGTAG and

JISS1R + NcoI AACT<u>CCATGG</u>TTGGGTTGTGTTTTATGTTGGTTTTG. The PCR fragment was cloned into pC1LUCP. Transgenic homozygous *JISS1:LUC* lines were generated in Col-5 background by floral dipping (92). Adapter ligation based PCR method was undertaken to identify the genomic location of *JISS1:LUC* insertion within At4g39240 (93).

Following *JISS1:LUC* crosses into mutants plants were determined as homozygous by diagnostic PCR for *JISS1:LUC* insertion using the following primers:

JISS1:LUC T-DNA F: GTCCTTGGTGGATGCATTGAT, JISS1:LUC T-DNA R: CTCCGTGCAACAGATTTTGGTT and JISS1:LUC T-DNA internal: GATCCCCCGAATTAATTCGGCG.

All other primers for validating mutants following crossing are shown in Table S2. All homozygous *JISS1:LUC* mutant crosses were generated in the mutant parental Col-0 background.

*JISS1* promoter: The JISS1^1-89^GFP line was generated using restriction enzyme cloning. The *JISS1* promoter and initial coding sequence was amplified from genomic DNA using primers;

JISS1:GFP pro FP: TCAAA<u>GAATTCC</u>GTAAAAGGTCGGTGTAGC and

JISS1:GFP RP: GTG<u>CCATGG</u>AGAAAAGCTCAGTTTCTGGATG. This PCR product was then cloned into the pCAMBIA1305 plasmid as a C-terminal fusion with GFP. The *JISS1* promoter full length *JISS1:GFP* line was generated using the Golden Gate assembly system (94) as follows; the *JISS1* promoter was amplified as above and cloned into the Level 0 plasmid pICH41295 while the full length *JISS1* coding sequence was cloned into pAGM41287. The *JISS1* promoter and coding sequence were combined in the FL1P1 level 1 vector to create ProJISS1:JISS1GFP:Ocs. Finally, the level 2 vector FL1P2 with BASTA selection was used to allow selection *in planta* after transformation into Col-0 background by standard methods.

Dex:*avrRpm1* plants were as described (34) and the *JAZ10:GUS* line was from (59).

### Bacterial growth, maintenance and inoculation

Bacterial cultures (*P. syringae* pv. *tomato* strains DC3000 [DC] containing the empty cloning vector (pVSP61), DC*hrpA* or DC3 containing the avirulence genes *avrRpm1, avrRps4* or *avrRpt2* were grown on Kings B medium (95) and prepared as described previously (90). *avrRpm1* (in pVSP61) was introduced into the DC coronatine-deficient mutant DB4G3 (*cor*^*-1*^*/cor*^*-2*^) (55). For luciferase, GUS and GFP assays, selected leaves were inoculated with a 1-ml needleless syringe on their abaxial surface with the appropriate bacterial suspension adjusted to a final optical density at 600 nm (OD_600_) of 0.15 in 10 mM MgCl_2_ (or as indicated in figure legends).

For SAR growth assays, the immunizing inoculation comprised either 10 mM MgCl_2_ (mock) or OD_600_ 0.005 DC3*avrRpm1* followed 2 days later by inoculation with *P. syringae* pv. *maculicola* strain M4 (*Psm4*) at OD_600_ 0.001 using a similar protocol to (96). Four days after *Psm4* challenge, bacterial growth measurements were determined from three challenged leaves per plant, and a minimum of four independent replicates. Significant growth differences between treatments were determined by Students t-test (P < 0.5), error bars representing the standard deviation (SD) of the mean. All experiments were repeated minimally three times.

### Chlorophyll fluorescence and biophoton imaging

Chlorophyll fluorescence measurement of challenged leaves were carried out using the parameters described (43) and *F*_*v*_*/F*_*m*_ data was extracted using the FluroImager Software. Biophoton imaging was carried out exactly as described (34).

### Luciferase visualisation

A solution of 1 mM luciferin (Promega) in 0.02% Silwet L77 (Loveland Industries, Ltd) was sprayed onto *JISS1:LUC* plants. The plants were kept in the dark for 30 min prior to inoculation. Petioles of the treated and adjacent rosette leaves were secured with minimise epi-nastic movement. Plants were placed in a dark box and images captured using either an ORCAII ER CCD camera (Hamamatus Photonics) with a 35 mm f2.8 micro Nikkor lens or Retiga R6 Scientific CCD camera (Qimaging) fitted with a Schneider STD XENON 25mm lens. Photons were counted for 10 min at 2 × 2 binning mode and data acquired with either Wasabi (Hamamatsu) or Micro-Manager 1.4 (Qimaging) software.

### Biophoton visualisation

Pathogen challenged plants were placed inside a dark box mounted with a Retiga R6 camera with 25mm f1.4 Navitar lens. Digital monochrome images captured photons for 20 mins at 2×2 binning mode over the appropriate time period using MicroManager v1.4. False colouring, brightness adjusting and annotation were performed using Fiji (97).

### Chemicals

JA, SA, ABA, azelaic acid (AZA), nonanoic acid (NA), pipecolic acid (Pip), DIECA (diethyldithiocarbamic acid), phenidone (1-Phenyl-3-pyrazolidinone), lanthanum (III) chloride (LaCl_3_) and coronatine were all from Sigma (Dorset, UK). N-hydroxyl-pipecolic acid (NHP) was synthesised by Accel Pharmtech (NJ 08816 USA) and Jarin-1 was sourced from (AOBIOUS Inc, USA). All chemicals were used at the concentrations described in the figures. Note, DIECA (2.5 mM), phenidone (2 mM), Jarin-1 (25 µM) or LaCl_3_ (1 mM) were either pre-infiltrated into systemic leaves, co-infiltrated with DC*avrRpm1* or the petiole of the local leaf was treated with the inhibitor soaked in cotton wool and secured in place with cling-film.

### JAZ10:GUS expression

GUS activity in systemic leaves was assessed at 4, 8 and 24 hpi by GUS staining (1 mM X-Gluc, 100 mM NaPO_4_ buffer pH 7.0, 10 mM EDTA, and 0.1% v/v Triton X-100). Samples were incubated at 37°C, and leaves were de-stained by repeated washes with 70% ethanol.

### Confocal imaging

Freshly excised leaf samples were mounted in water and imaged on a Zeiss LSM 880 confocal microscope with a 100× oil immersion (ER images) or 10x air objective (whole leaf images) lens. GFP was excited at 488nm and detected in the 498-559nm range; chlorophyll A was excited at 561nm and detected in the 605-661 nm range. All image analysis was performed in Fiji (97)

### Electrophysiology experiments

Surface electrical potentials of challenged and systemic leaves were measured following infection using four electrodes (Fig. 5A) adopting a similar approach to (59). The working electrode (W1, red) was always placed on the laminar immediately above the petiole of the challenged leaf. Similarly, electrodes W2 (green) and W3 (blue) were placed on adjacent systemic leaves. Electrode W4 (black) was placed on the distal systemic leaf. The reference electrode (Ref) was placed in the soil. Surface potential changes were measured from the reference electrode to the working electrode. Electrical recordings were captured using a data logger (PicoLog 1000) and signal amplitude and duration were plotted for each leaf over ∼24h. Control recordings over an extended time period predominately showed a constant surface potential.

## Acknowledgments and funding sources

We thank Rebecca Winsbury (Exeter) and Nestoras Kargios (Warwick) for technical help and Peter Winlove and Stephen Green (Exeter) for electrophysiology advice. Xinnian Dong (UC Duke) and Edward Farmer (University Lausanne) provided *Arabidopsis* mutants used in this study. Craig Gall helped with figure preparations.

## Author contributions

MG, MTZ, TG, SB and EB conceptualized and designed the experiments. TG, SB, EB and RH performed the experiments and analysed data. TG, MTZ, RH and SK generated material. DH designed equipment, and LF and DH provided experimental insights. MG, SB, EB prepared the manuscript. EB, SB and MG wrote the manuscript.

## Data and materials availability

All data is available in the main text or the supplementary materials. *Arabidopsis thaliana* reporter lines are available from the corresponding author.

