## Supplemental Figures and Tables for "Rapid local and systemic jasmonate signalling drives initiation and establishment of plant systemic immunity"

### Supplementary Figures

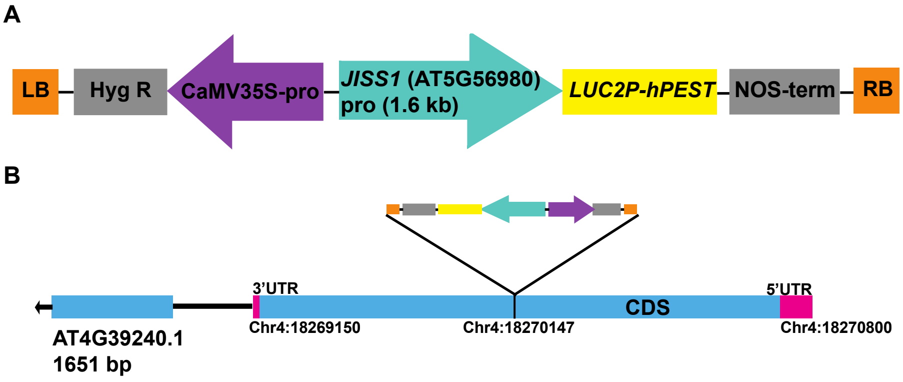


**Figure S1: Schematic of the *JISS1:LUC* construct and its T-DNA integration location in the *Arabidopsis* genome.** The *Photinus pyralis LUC2P* reporter gene containing the hPEST protein destabilization sequence (reduces intracellular half-life) was PCR amplified and cloned into pCAMBIA1302 digested with Kpn1 and Pml1 to generate pC1LUCP with a *NOS* terminator. The 1631 bp *JISS1* promoter was PCR amplified from Col-0 genomic DNA using the primers as described (1) generating a hygromycin selectable *JISS:LUC* construct **(A)**. **(B)** The *JISS1:LUC* construct was agro-infiltrated into *A. thaliana* ecotype, Col-5 and stable homozygous lines were selected. Subsequent sequence analysis as per (2) identified the genomic insertion point of the *JISS1:LUC* T-DNA construct at position 18270147 in the coding region of AT4G39240. Eight independent lines were generated and tested for expression in systemic leaves following an immunising challenge with DC*avrRpm1*. This line was used for all further studies and crosses.

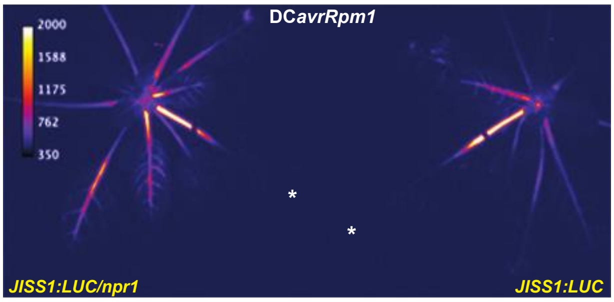


**Figure S2: *JISS1* signal propagation is not impaired in known SA signalling mutants.** An *JISS1:LUC/npr1* line exhibited comparable systemic luciferase activity to that of *JISS1:LUC* (right) following challenge with DC*avrRpm1* (4:20 hpi). A white asterisk denotes treated leaves. Images are false coloured by signal intensity, as indicated by calibration bar.

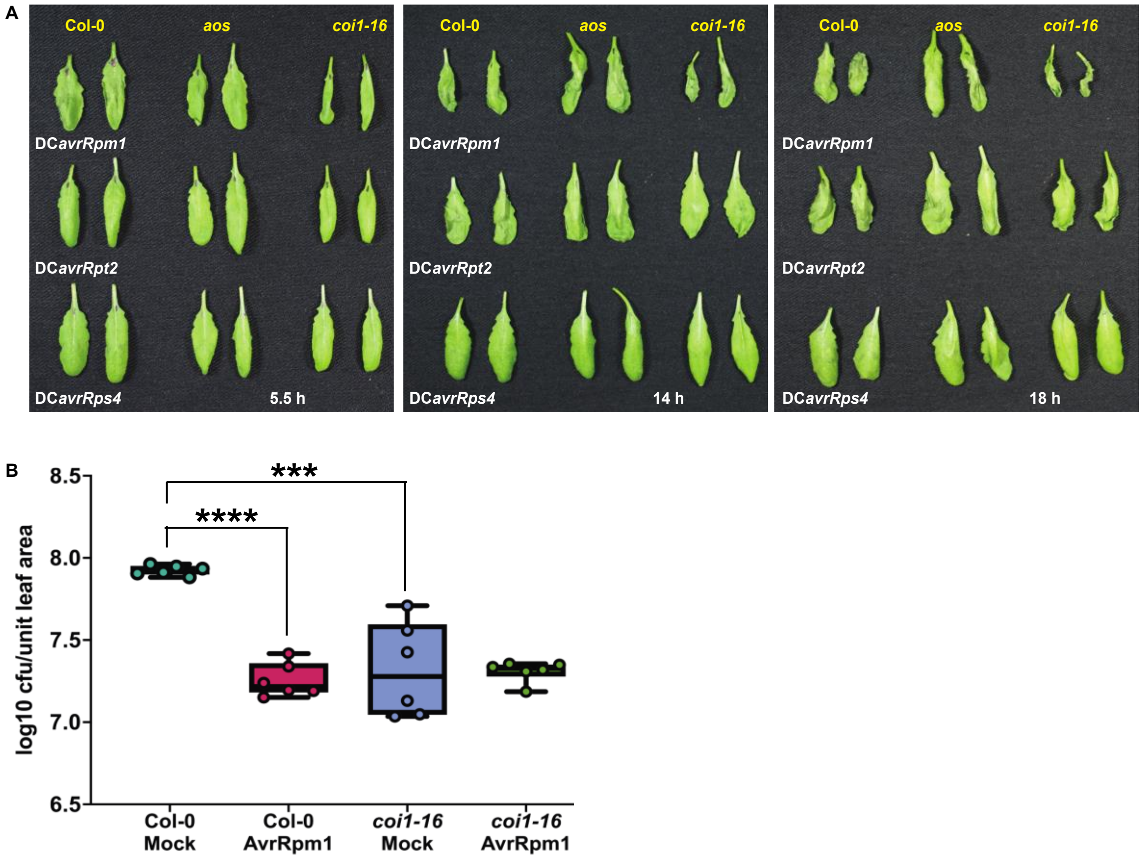


**Figure S3: HR development is not affected in mutants with impaired JA biosynthesis (*aos*) or perception (*coi1-16*) but *coi1-16* has reduced disease susceptibility. (A)** Phenotyping of *aos* and *coi1-16* mutants show comparable HR development to Col-0 plants, indicating that loss of JA biosynthesis and detection does not affect HR development. **(B)** SAR growth curve of *Psm4* following DC*avrRpm1* or mock pre-treatment on wild-type (Col-0) and the *coi1-16* mutant. Error bars represent the mean ± SE (n=6) and statistical significance (P<0.0001) was determined by Student t-test. This is representative of 3 independent experiments.

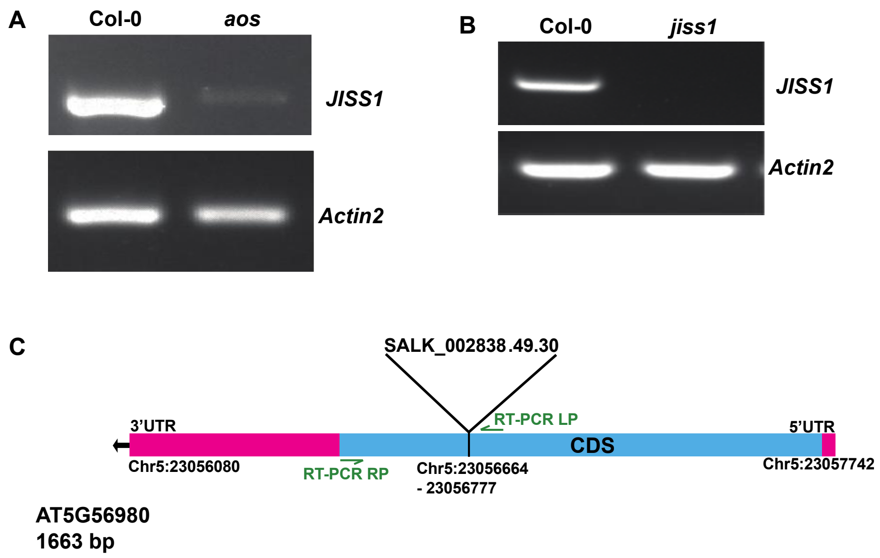


**Figure S4: *JISS1* expression is absent in the systemic leaves of the JA biosynthetic mutant, *aos* and *jiss1* mutant. (A)** Representative RT-PCR analysis of *JISS1* expression in systemic leaves from *DCavrRpm1* challenged Col-0 and the JA deficient *aos* mutant demonstrates *JISS1* induction is dependent on *de novo* JA synthesis. *Actin2* expression was used as a loading control. **(B)** RT-PCR verification that *JISS1* expression is knocked out in the *jiss1* T-DNA insertion line. *Actin2* expression was used as a loading control (3). **(C)** T-DNA insertion in the coding sequence of the *JISS1* gene (AT5G56980) is located at position Chr5:23056664 – 23056777. Relative positions of forward and reverse gene specific RT-PCR primers are indicated and sequences are given in Methods.

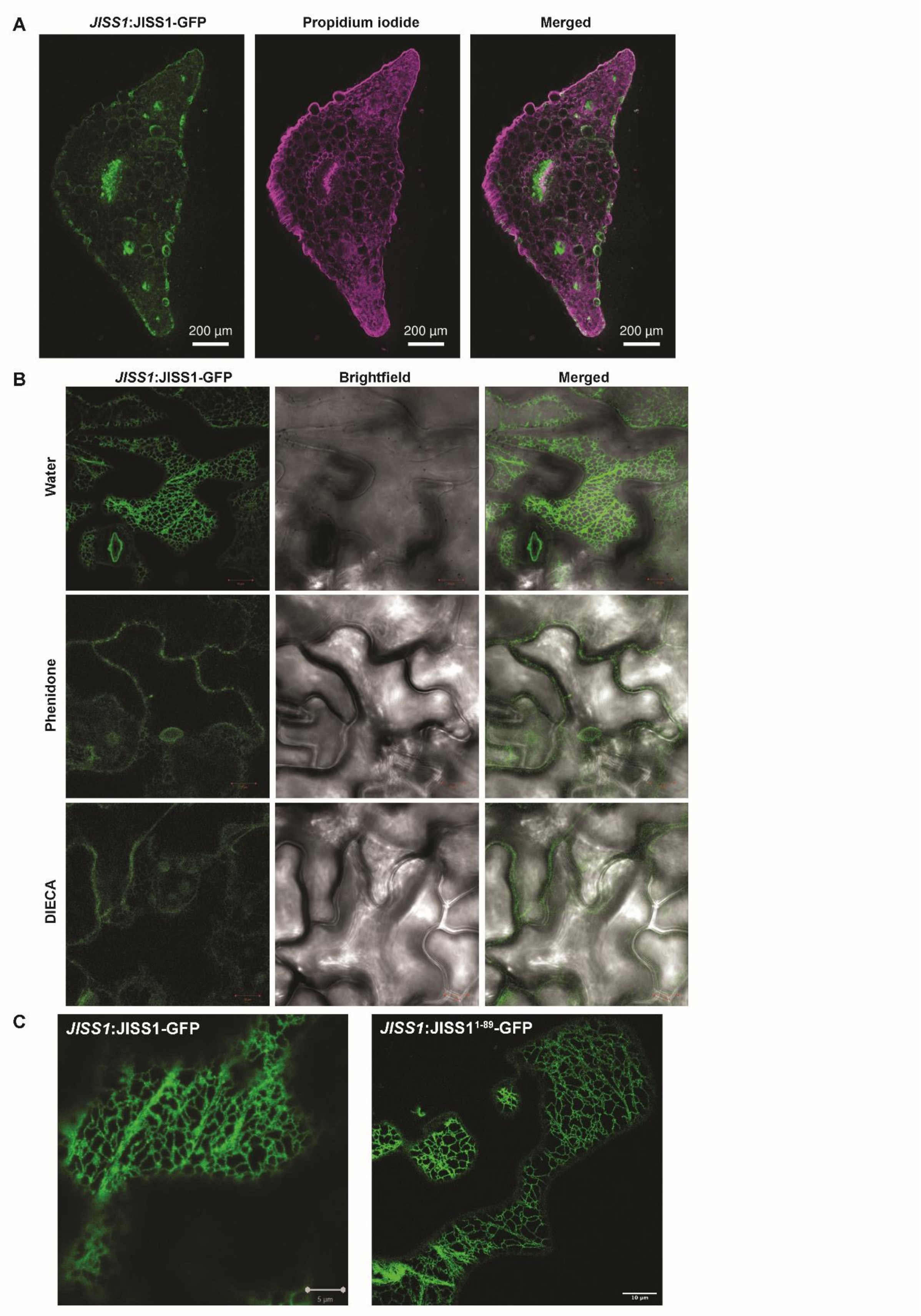


**Figure S5: JA dependent localisation of *JISS1:*JISS1$^{1\text{-}89}$*-GFP* and *JISS1:*JISS1*-GFP* in systemic tissue. (A)** Petiole section from systemic leaf showing propidium iodide (magenta) and *JISS1:*JISS1$^{1\text{-}89}$*-GFP*

expression in green and a merged image. Scale bar, 200 µm **(B)** *JISS1:*JISS1$^{1-89}$-GFP systemic leaves pre-treated with JA inhibitors, phenidone and DIECA prior to DC*avrRpm1* challenge*.* Mock treatment of systemic leaves shows JISS1$^{1-89}$-GFP labelling of the entire ER network 4 hpi whereas systemic leaves infiltrated with phenidone or DIECA exhibit minimal GFP signal in the ER. **(C)** *JISS1:*JISS1-GFP and *JISS1:*JISS1$^{1-89}$-GFP plants following DC*avrRpm1* challenge, showing comparable subcellular localisation of GFP in endoplasmic reticulum of epidermal cells. Scale bars, 5 µm and 10 µm, respectively. All confocal images are representative of multiple cells from multiple experiments.

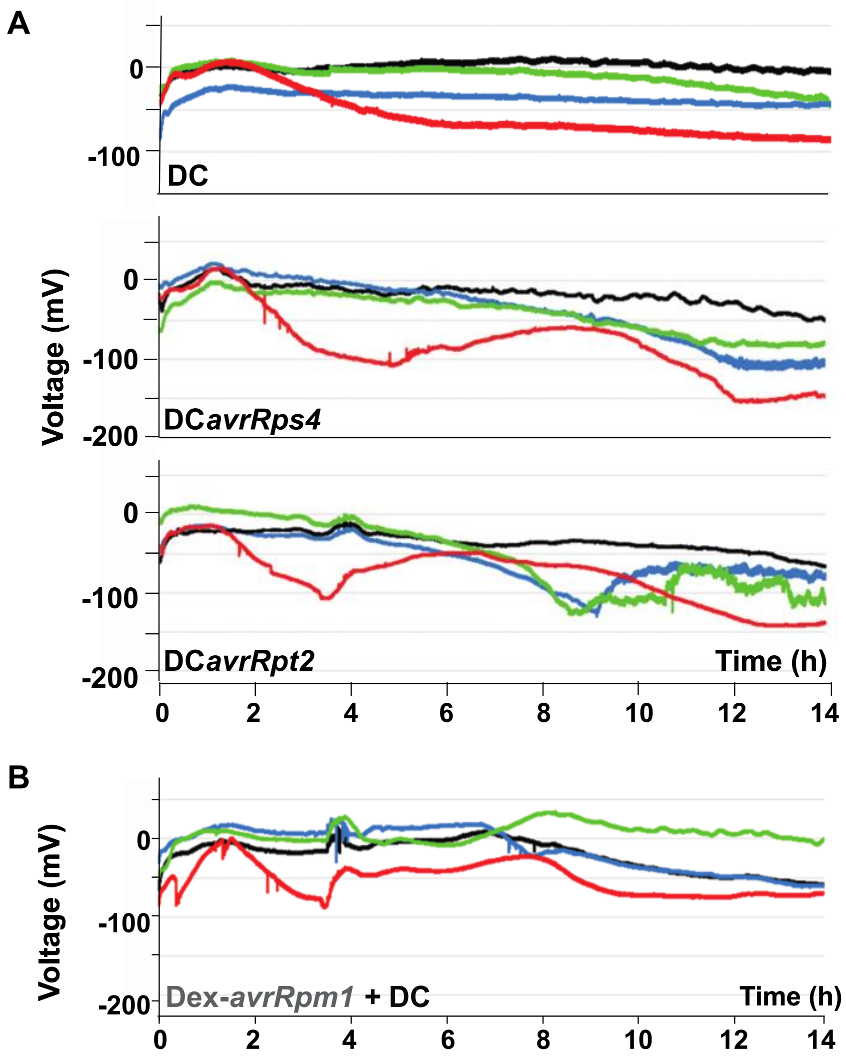


**Figure S6: ETI activation of systemic electric signalling. (A)** Col-0 plants do not show development of SISPs following DC challenge (top trace) whereas challenge with DC*avrRps4* (middle trace) or DC*avrRpt2* (bottom trace) exhibits depolarisation of the infected leaf from 2 hpi, followed by induction of SISPs from 6-10 hpi. **(B)** Infiltration of DC 1 h after Dex induction of *avrRpm1* induces depolarisation and repolarisation of the infiltrated leaf and initiation of SISPs, albeit weaker than DC*avrRpm1*.

**Table S1**

| **AGI** | **Annotation** | **Mean fold change (DC*avrRpm1*/DC*hrpA*)** |
|---|---|---|
| At1g19180 | *JAZ1* | 4.55 |
| At1g74950 | *JAZ2* | 3.35 |
| At3g17860 | *JAZ3* | 1.86 |
| At1g17380 | *JAZ5* | 8.15 |
| At1g72450 | *JAZ6* | 3.41 |
| At2g34600 | *JAZ7* | 5.47 |
| At1g30135 | *JAZ8* | 5.88 |
| At1g70700 | *JAZ9* | 3.49 |
| At5g13220 | *JAZ10* | 6.59 |
| At1g48500 | *JAZ4* | No probeset |
| At3g43440 | *JAZ11* | No probeset |
| At5g20900 | *JAZ12* | Not differentially expressed |
| At5g56980 | *JISS1* | 4.20 |

JAZ genes significantly induced at 4 hpi with DC*avrRpm1* compared to DC/DC*hrpA* challenges, together with *JISS1* expression data in all treatments. Mean fold change ratios derived from normalised expression values on ATH1–121501 Affymetrix GeneChips as described (4). The data set is deposited at http://affymetrix.arabidopsis.info/narrays under identifier NASCARRAYS-403.

**Table S2: Primers for identification of homozygous T-DNA lines after crosses**.

* These lines were derived from the *npr1/3/4* triple mutant obtained from the lab of Xinnian Dong (5).

| **Mutant** | **Gene** | **t- DNA** | **Primers Sequence (5' to 3')** | **KO. Product size** |
|---|---|---|---|---|
| ***sid2*** | AT1G74710 | SALK_133146.39.30 | LP - TCTGATGGATCTCCAATCGTC<br>RP - GAGATTTCAAGACGCCACTTG | 577-877 |
| ***nac055*** | AT3G15500 | SALK_014331.54 | LP - TAAACGATGAGCGATAGCGAG<br>RP - AAAGGAACCAAAACCAATTGG | 467-767 |
| ***nac019*** | AT1G52890 | SALK_096295.49.30 | LP - TCAATGAACTCAAGGGATTGC<br>RP - ATGCGGTTTGGGTTAGAAAAC | 459-759 |
| ***nac072*** | AT4G27410.2 | SALK_083756.50.50 | LP - GACTGGTCTTTTATCTCCGGG<br>RP - ACAACACATCGATAAGGTCGG | 527-827 |
| ***jiss1*** | AT5G56980 | SALK_002838.49.30 | LP-ATGTTTACCCGGATCCAAATC<br>RP-GCCACACATACTTCGCTAAGC | 552-852 |
| ***coi1-16*** | AT2G39940 | SALK_045434 | | |
| ***fmo1*** | AT1G19250 | SALK_026163 | LP- CTTTTCGGTTGGACTTGGAAC<br>RP- CTGCTTTGGACGTATCCTACG | 485-785 |
| ***aos*** | AT5G42650 | SALK_017756 | LP- CGAGAAATTAACGGAGCTTCC<br>RP-CTAACCGGAGGCTACCGTATC | 432-732 |
| ***glr3.3a*** | AT1G42540 | SALK-099757 | LP-GATGCTGCATATGGTTGTGTG<br>RP-GTTGAACGATAAGCTTGCGAG | 700 |
| ***glr3.6a*** | AT3G51480 | SALK_091801 | LP-TTCGTTCAAAGGTGGCATAAC<br>RP-CGACTATGAGGAAAGACGCAG | 550 |
| ***npr3* (deleted)*** | AT5G45110 | SALK_043055 | LP1-TGATTGTTGTCGACCTGCCA<br>RP1-AGATCTGACCTCGCCACTCT<br>LP2-TTGGTTCTTTTGCCTTCTCTTTGA<br>RP2-GGCATCCCTATCACCATCTGT | 307<br><br>209 |
| ***npr4**** **(deleted)** | AT4G19650 | SALK_098460 | LP1-TTGGCGATGAAGCTAAGGGG<br>RP1-CTGGCAGAGAGCATGAACCA<br>LP2- TACGCTACTGCTGTTCCAGA<br>RP2-CTTGCACGTGTGCTTTTTGG | 526<br><br>341 |
| ***npr1**** | AT1G64280 | EMS mutagenized | LP-CTCGAATGTACATAAGGCAC<br>RP-GTGCGGTTCTACCTTCC | 296 |

**Movie S1**

***JISS1* expression is induced systemically by ETI.** Temporal spatial dynamics of luciferase activity in *JISS:LUC* plants following DC*avrRpm1* (4 hpi, bottom left and top right plant), DC*avrRps4* (13:20 hpi, top left and top right plant) and DC*avrRpt2* (15:20 hpi, bottom right and top right plant). White asterisk indicates infiltrated leaves, movie is false coloured by signal intensity.

**Movie S2.**

**JISS1 signal propagates symplastically through the epidermis and vascular tissue.** JISS1$^{1\text{-}89}$-GFP expression is predominately restricted to the central vein and epidermal cell layer.
